# Prostate Specific Membrane Antigen expression in the vasculature of primary lung carcinomas associates with faster metastatic dissemination to brain

**DOI:** 10.1101/863456

**Authors:** Jayendrakishore Tanjore Ramanathan, Suvi Lehtipuro, Harri Sihto, József Tóvári, Lilla Reiniger, Vanda Téglási, Judit Moldvay, Matti Nykter, Hannu Haapasalo, Vadim Le Joncour, Pirjo Laakkonen

**Author notes:** Corresponding author: Prof. Pirjo Laakkonen, Translational Cancer Medicine Research Program, Haartmaninkatu 8, 00014 University of Helsinki, Finland.

## Abstract

Glioblastomas and brain metastases (BM) of solid tumors are the most common central nervous system neoplasms associated with very unfavorable prognosis. In this study, we report the association of Prostate Specific Membrane Antigen with various clinical parameters in a large cohort of primary and secondary brain tumors. A tissue micro array containing 371 cases of ascending grade gliomas pertaining to astrocytic origin, samples of 52 cases of primary lung carcinomas with matching brain metastases with follow-up time accounting to 10.4 years was evaluated for PSMA expression using immunohistochemistry. In addition, PSMA expression was studied in brain metastases arising from melanomas and breast carcinomas. Neovascular expression of PSMA was evident alongside with high expression in the proliferating microvasculature of glioblastomas when compared to the tumor cell expression. This result corroborated with the results obtained from the *in silico* (cancer genome databases) analyses. In the matched primary lung cancers and their brain metastases (n = 52), vascular PSMA expression in primary tumors led to significantly accelerated metastatic dissemination to the brain with a tendency towards poor overall survival. Taken together, we report the vascular expression of PSMA in the primary and secondary brain tumors that globally associates with the malignant progression and poor outcome of the patients.

## 1. Introduction

Gliomas, neoplasms of glial cell origin, constitute approximately 30% of all nervous system tumors and 80% of all malignant brain tumors (1). Glioblastoma, a grade IV astrocytoma, is the most frequent, aggressive and lethal glioma (2). Glioblastoma harbors a dense abnormal vasculature, display large hypoxic and necrotic areas and contain extensively proliferating tumor cells with the intrinsic ability to disseminate and colonize the organ far beyond the principal tumor mass. The current standard of care comprising surgery followed by radio- and chemotherapy, provides only a modest improvement in the progression-free and overall survival, and patient prognosis remains dismal (3, 4). Brain metastases (BM) of solid tumors are the most frequent intracranial tumors and about 10-fold more common than primary brain tumors (5). Brain metastases are associated with severe neurological symptoms and abysmal prognosis compared with other metastatic sites (6). The primary tumors most often responsible for brain metastases are melanomas (5-20%), lung (36-64%), and breast cancers (15-25%) (7).

Prostate Specific Membrane Antigen (hereon referred to as PSMA) also known as Glutamate Carboxypeptidase II (GCP II), N-acetyl-L-Aspartyl-L-glutamate peptidase I (NAALDase I) or N-acetyl-aspartyl-glutamate (NAAG) peptidase is a transmembrane glycoprotein encoded by the *FOLH1* gene. Until today, only two well-defined physiological roles of PSMA are known; the folate-hydrolyzing activity in the small intestine and the NAAG-hydrolyzing activity in the nervous system (8, 9). PSMA is expressed in tumor cells of almost all prostate cancers and its increased expression is associated with tumor aggressiveness, metastasis, and recurrence (10, 11). Numerous studies have shown that the normal vasculature is devoid of PSMA expression while tumor neovasculature often shows high PSMA expression (12-14). Moreover, PSMA expression has been reported in benign gliomas (grade I), malignant glioblastomas (grade IV), and breast cancer brain metastases with very limited number of patients (15). Conway *et al.* 2006 (16) showed that PSMA expression is critical during angiogenesis as PSMA null mice were unable to mount a pathological angiogenic response (16). In addition, Grant *et al.*, 2012 reported that PSMA-regulated angiogenesis was independent of VEGF by using a mouse model of oxygen-induced retinopathy (17).

Our study is in line with the previous reports about PSMA expression in the tumor neovasculature. However, we accompany our immunohistochemical study with the PSMA mRNA level expression analyses in large genetic datasets. We report higher PSMA expression in glioblastomas compared to lower-grade gliomas. For the first time, we report PSMA expression in the proliferating microvasculature of glioblastomas, which is one of the histopathological features that differentiates glioblastoma from the lower-grade gliomas. We also had an exceptional opportunity to screen rarely available specimens of the matched primary lung cancers and their brain metastases as well as samples from melanoma and breast brain metastases. Importantly, PSMA expression in the primary lung tumor vasculature associated with significantly accelerated metastatic dissemination to the brain with a tendency towards poor overall survival. Overall, the large sample size along with the *in silico* data analyses provide more comprehensive view of the role of PSMA compared with prior studies involving much smaller patient cohorts.

## 2. Materials and Methods

### 2.1 Patient material

#### i) Gliomas

The ethical committee of Tampere University Hospital and the National Authority for Medico-legal Affairs in Finland approved the usage of tumor tissue array and the study design. The tumor samples were collected from the patients who underwent surgery at the Tampere University Hospital during the years 1983-2009. The astrocytoma specimens were initially fixed in 4% phosphate-buffered formaldehyde and then processed into paraffin blocks. One histologically representative tumor region was selected from each specimen. From the selected regions, 0.6 mm tissue cores were mounted into tissue microarray (TMA) blocks (18). Our tumor series included 371 gliomas of astrocytic origin ranging from 41 (11.05%) grade I pilocytic astrocytomas, 46 (12.39%) grade II diffuse astrocytomas, 25 (6.73%) grade III anaplastic astrocytomas, and 259 (69.81%) grade IV glioblastomas (Table 1) based on the diagnosis at the time of the surgery. Of these 371 gliomas, 288 were primary and 83 recurrent tumors from 174 men and 114 women (Table 1).

**Table 1.** Clinicopathologic features of the astrocytomas.

| **Astrocytoma** | **Grade I** | **Grade II** | **Grade III** | **Grade IV** | **Total** |
| --- | --- | --- | --- | --- | --- |
| *No. of patients*(†) | 41 | 46 | 25 | 259 | 371 |
| **Primary** | 33 | 34 | 16 | 205 | 288 |
| *Residual* |  |  |  |  |  |
| **Second** | 7 | 11 | 6 | 43 | 67 |
| **Third- sixth** | 1 | 1 | 3 | 11 | 16 |
| *Sex (n)* (§) |  |  |  |  |  |
| **Male** | 20 | 24 | 8 | 122 | 174 |
| **Female** | 13 | 10 | 8 | 83 | 114 |
| *Age (y)* (§) |  |  |  |  |  |
| **Median** | 11 | 40 | 39.50 | 62.00 | - |
| **Mean ± SD** | 18.4 ± 16.7 | 42.8 ±17.5 | 42.2±14.5 | 61.4±11.0 | - |
| Minimum | 1 | 12 | 23 | 4 | - |
| Maximum | 57 | 83 | 81 | 84 | - |
| *Follow-up* |  |  |  |  |  |
| **End Survivors (n)** | 26 | 13 | 5 | 7 | 51 |
| **5-year survival (%)** | 81,3 | 38,2 | 31,3 | 3,4 | 17,8 |

(†) includes primary and secondary tumors
(§) includes only primary tumors

#### ii) Primary lung tumors and brain metastases

Samples of surgically resected, paired primary lung carcinomas and their associated brain metastases from 52 patients, as well as breast carcinoma BMs from 18 patients and melanoma BMs from 19 patients were analyzed in this study. Additionally, TMAs consisted of primary lung carcinomas from 3 patients and lung BMs from 4 patients apart from the paired samples. Permissions to use the archived tissues have been obtained from the Local Ethical Committee of the Semmelweis University (Budapest, Hungary) (TUKEB-1552012, −5102013 and −862015) and the study was performed in compliance with the Declaration of Helsinki. The samples were fixed in 10% neutral-buffered formalin, dehydrated and cleared in increasing concentration of ethanol and xylol before embedding into IHC-grade paraffin. For the tissue microarray (TMA), hematoxylin and eosin-stained sections were used to define the tumor areas and three representative 2 mm cores were obtained from each case and inserted in a grid pattern into a recipient paraffin block using a tissue arrayer (3D Histech, Budapest, Hungary).

### 2.2 Immunohistochemistry

The PSMA and CD31 expression was detected from the sections by using the TSA kit (Perkin Elmer) according to the manufacturer’s instructions. Tumor samples were deparaffinized by immersing the slides in xylene and rehydrated with graded ethanol (100%-70%) and water. Briefly, the antigen was retrieved with heat treatment in citrate buffer (1.8 mM citric acid, 8.2 mM sodium citrate, pH 5.0). The endogenous peroxidase activity was irreversibly inactivated by incubating the sections with 3% H2O2-MetOH for 10 minutes and blocked with TNB buffer (0.1M Tris-HCl (pH 7.5), 0.15M NaCl, 0.5% blocking reagent from TSA kit) for 30 minutes. The primary antibody, rabbit-anti-human mAb for PSMA (Abcam, 4 µg/ml) and mouse anti-human CD31 (DAKO) was applied overnight at 4°C. Post primary antibody incubation and washes with TNT (Tris/NaCl/Tween 20) buffer, the sections were incubated with the biotinylated secondary antibody (goat anti-rabbit and goat anti-mouse, Dako) for 60 minutes. The staining was amplified by using the SA-HRP and biotinylated tyramide and finally visualized using the AEC (0.95M 3-amino-9-ethylcarbazole, 6% N,N dimethylformamide, 94% Sodium Acetate, 0.01% H2O2). The sections were counterstained with Mayer’s hematoxylin.

### 2.3 Evaluation of immunostaining and statistical analysis

Overall analysis was performed for all patients whose tumor tissue was available for immunohistochemistry. PSMA stained sections were scanned using the 3DHistech Panoramic 250 FLASH II digital slide scanner at an absolute magnification of 20×. The evaluation of the scanned sections was performed blinded to the clinical data. The PSMA staining in the TMAs was scored as negative (0) or positive (1) (Figure S1). The samples in the TMA that lacked vascular structures were marked as empty and excluded from the analysis. We graded separately the vascular and cellular expression of PSMA and associated it with the clinical and molecular pathological parameters. Overall survival was calculated from the date of primary tumor diagnosis to date of death censoring the patients who were alive at the end of the follow-up period and the progression-free survival from the date of diagnosis to date of tumor progression or death, censoring patients without detected progression or death. Brain metastasis-free survival was calculated from the date of primary tumor diagnosis to date of brain surgery. Brain metastasis evolved in all patients in the cohort. The statistical analysis was performed using the IBM SPSS statistics 21.0 software for Windows. Chi-square and non-parametric tests were used to analyze the associations between PSMA expression and various clinical and molecular pathological parameters. Overall, progression-free and brain metastasis-free survival were analyzed using the Kaplan-Meier and survival between the groups were compared using the Log-rank and Breslow tests. The overall survival in primary lung carcinomas was analyzed in 55 patients including both the paired (n = 52) and unpaired (n = 3) samples. The analysis of time to metastatic dissemination was performed using only the paired (n = 52) patient samples. A *P*-value of < 0.05 was considered significant. Statistical analyses for public datasets were performed using the R version 3.2.2.

### 2.4 Tumor datasets

The clinical data and level 3 mRNA microarray data (Agilent 244K TCGA Custom 1-2) of primary GBM samples were downloaded from the NCI Genomic Data Commons. Combining multiple samples from the same patient using median and removing samples without clinical data resulted in 567 samples. FPKM values for the GBM anatomic structures (122 samples) and sample information were downloaded from the Ivy GAP website (http://glioblastoma.alleninstitute.org/) (19). Data were log2 transformed. We have used the nomenclature based on that of the TCGA.

## 3. Results

### 3.1 PSMA is highly expressed in the hyperplastic proliferating microvasculature of grade IV gliomas

To evaluate the PSMA protein expression in human glioma samples, TMAs containing 371 grade I-IV gliomas were stained using the anti-PSMA antibody. Vascular expression of PSMA (Figure 1A). was validated by visualizing the blood vessels using an anti-CD31 antibody (Figure 1B). We graded PSMA expression as positive (1) or negative (0). For the vascular expression, we excluded the samples that did not show any vascular structures in the TMA-samples. PSMA expression was most frequently detected in the vasculature of glioblastomas (30.8%, 74/166) (*P* = 0.044) (Table 2) highlighting the distinct angiogenic feature of glioblastomas. The less angiogenic, lower-grade gliomas showed reduced PSMA expression in their vasculature: grade III (14.2%, 3/18), grade II (17.9%, 7/32), and grade I (15%, 6/34) (Table 2). We next analyzed PSMA expression in tumor cells to address whether an association exists between the vascular and cellular expression of PSMA. About 1/3 of the grade I (31.7%, 13/28) and II tumors (32.6%, 15/31) showed PSMA expression in tumor cells whereas (56%, 14/11) of grade III and (44.7%, 116/143) of grade IV gliomas (Table 2) expressed PSMA in tumor cells. However, the overall expression in tumor cells was lower than the one detected in the tumor blood vessels.

**Table 2.** Histological classification of astrocytic tumors based on the PSMA expression.

| *Tumor* |  | *Pilocytic* | *Diffuse* | *Anaplastic* | *Glioblastoma* | *Total* | *P – Value* |
| --- | --- | --- | --- | --- | --- | --- | --- |
| *grade* |  | *Grade I* | *Grade II* | *Grade III* | *Grade IV* |  | *Chi-square* |
| **PSMA (+/-)** | Vascular | 15% | 17.9% | 14.2% | 30.8% | 26.47% | **0.044** |
|  |  | (6/34) | (7/32) | (3/18) | (74/166) | (90/250) |  |
| **PSMA (+/-)** | Tumor | 31.7% | 32.6% | 56% | 44.7% | 42.5% | 0.102 |
|  | cells | (13/28) | (15/31) | (14/11) | (116/143) | (158/213) |  |

**Figure 1.**
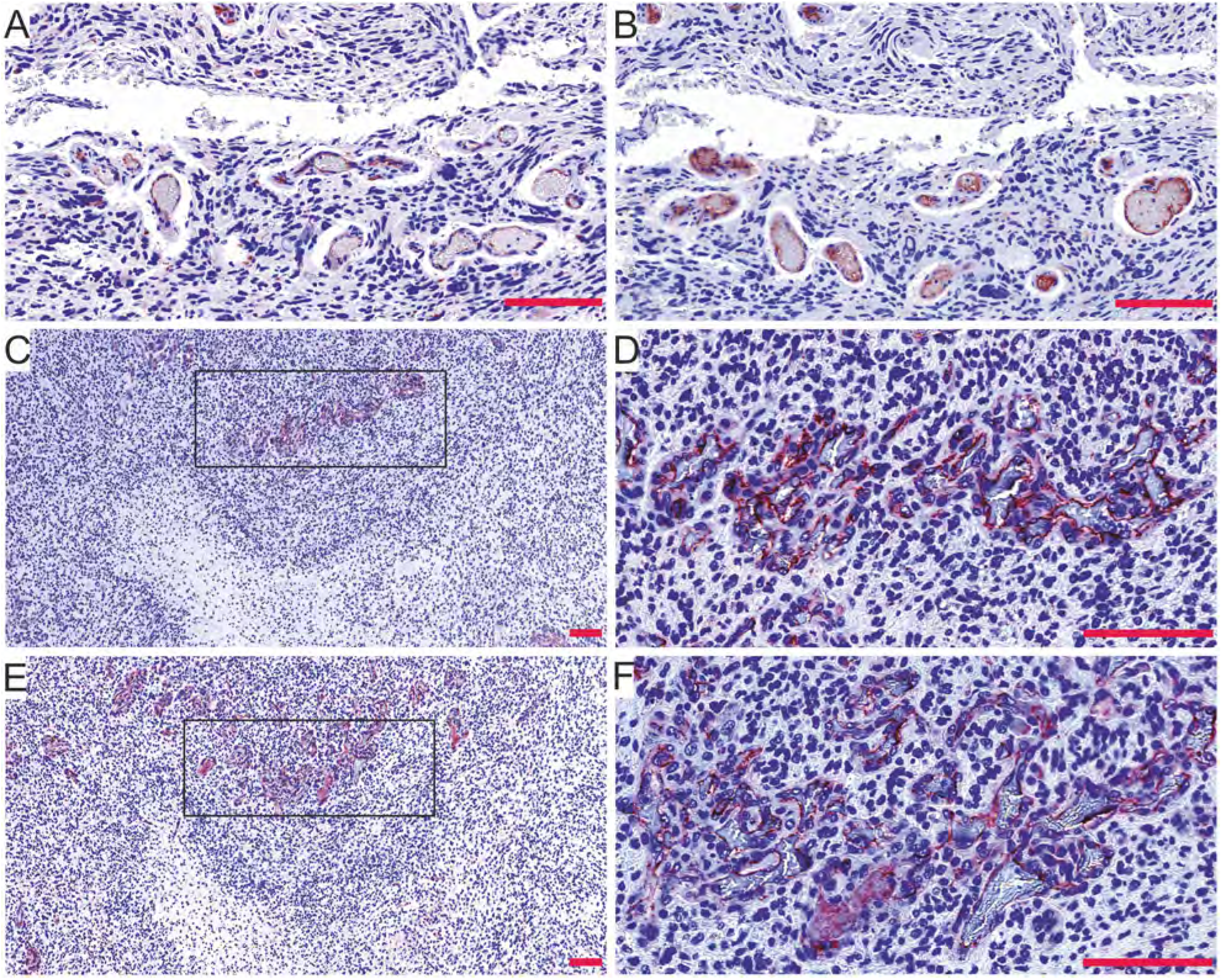
Expression of the Prostate Specific Membrane Antigen (PSMA) in gliomas. A. PSMA expression in the vasculature of grade IV glioblastoma was validated by staining the blood vessels using anti-CD31 antibody (B) in a subsequent tissue section (20x). C-F. The PSMA (C,D) was highly expressed by the hyperplastic proliferating microvasculature visualized by CD31 staining (E,F) Scale bar = 100 μm.

One of the diagnostic histopathological features that differentiates glioblastoma from the lower-grade gliomas is the presence of proliferating hyperplastic microvasculature that appears as tufted glomeruloid bodies and exist adjacent to the necrotic foci with bordering pseudopalisading cells. The glomeruloid tuft-like structures in certain tumor samples in the TMAs were positive for PSMA. Therefore, we analyzed the entire tumor block of two grade IV glioblastomas that were highly positive for the vascular expression of PSMA. This analysis confirmed that not the pre-existing mature vessels, but the proliferating hyperplastic microvasculature expressed high levels of both PSMA (Figure 1C,D) and a vascular marker CD31 (Figure 1E,F). To study PSMA (*FOLH1*) expression in the glioblastoma subtypes, we analyzed 567 mRNA microarray samples using the TCGA GBM dataset (see Materials and Methods, nomenclature from the TCGA). When we compared the expression of PSMA between the different subtypes, we observed decreased PSMA expression in the classical (CL) subtype compared to the other subtypes (*P* < 0.01, Wilcoxon rank-sum test) (Figure 2A).

**Figure 2.**
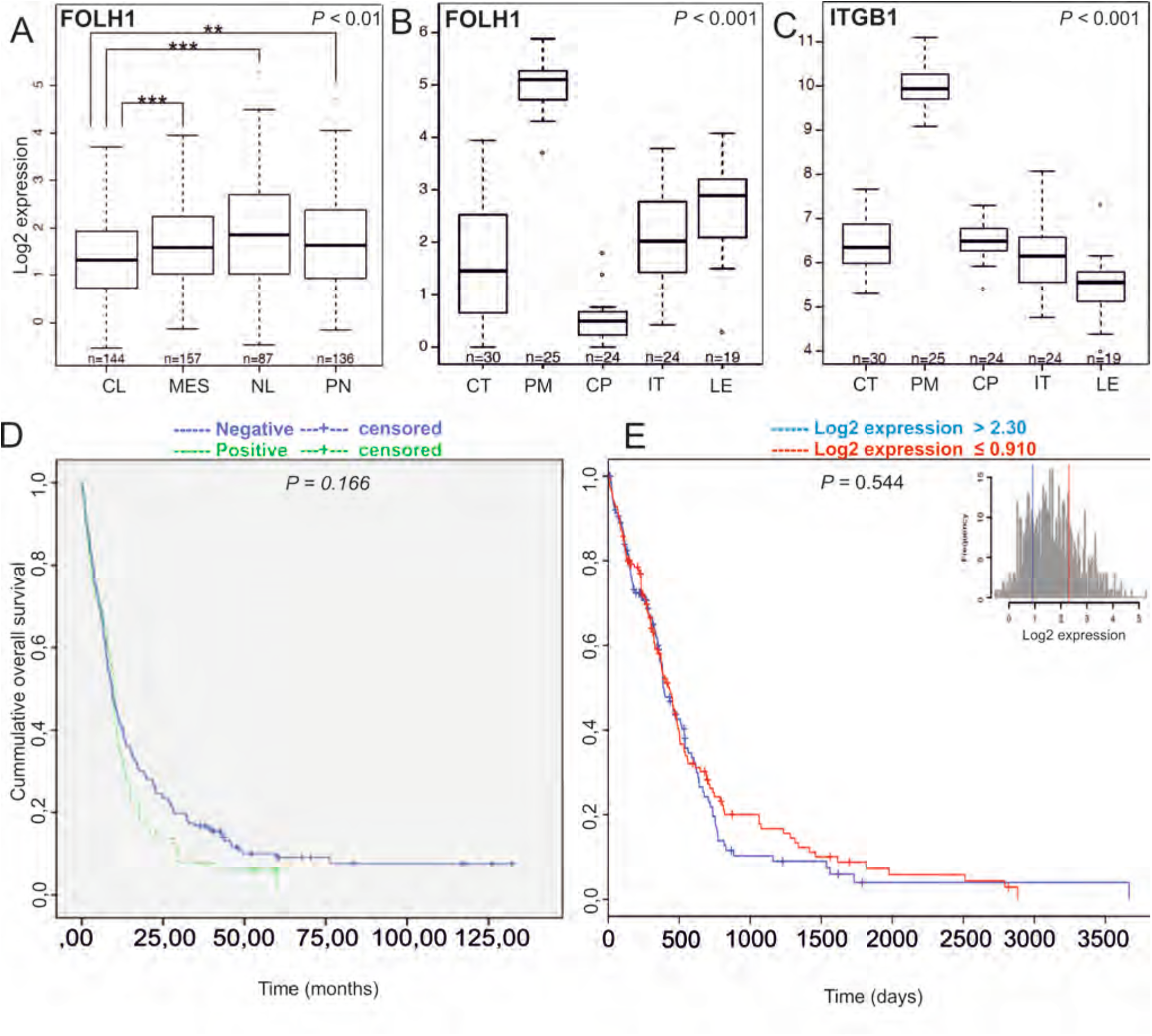
Analysis of PSMA/*FOLH1* mRNA expression using the TCGA database and association of PSMA/*FOLH1* expression with patient survival. A. Expression of PSMA/*FOLH1* was lower in the classical subtype (CL) compare to the other subtypes. *P* < 0.01**. B.** The proliferating microvasculature (PM) showed the highest and the cellular pseudopalisades (CP) the lowest PSMA expression when compared to other groups *P* < 0.001.C. Integrin β1 is specifically upregulated in the proliferating microvasculature (PM) similar to the PSMA/*FOLH1* expression. D-E. There was no difference in the overall survival of the patients based on the PSMA expression at the protein level (D, patient samples: n = 287, *P* = 0.166) or mRNA level (E, TCGA samples: n = 567, *P* = 0.544).

Next, we analyzed PSMA (*FOLH1*) expression using the IvyGAP RNA seq dataset (http://glioblastoma.alleninstitute.org/) that in contrast to other cancer genome datasets, shows the individual gene expression profiles for the different anatomic structures (such as leading edge (LE), infiltrating tumor (IT), cellular tumor (CT), proliferating microvasculature (PM), and cellular pseudopalisades (CP), which were isolated by using the laser microdissection (19). Consistent with our findings, proliferating microvasculature (PM) showed the highest (*P* < 0.001, Wilcoxon rank-sum test) and the cellular pseudopalisades (CP) the lowest PSMA expression when compared to other groups (*P* < 0.001, Wilcoxon rank-sum test) (Figure 2B). Since it has been reported that PSMA is involved in angiogenesis by activating the laminin-mediated integrin β1 (ITGB1) signaling (16), we analyzed the integrin β1 expression using the IvyGAP. Also, in this dataset, integrin β1 (*ITGB1*) was upregulated in the proliferating microvasculature (PM) when compared with all the other anatomical structures (Figure 2C) (*P* < 0.001).

### 3.2 PSMA did not associate with patient prognosis in gliomas

We next analyzed the association of PSMA protein level with patient survival in our TMA-material using the log-rank test. Neither the presence nor the absence of PSMA expression in the vasculature associated with the overall survival of patients (n = 287) (*P* **=** 0.166, log-rank test (Figure 2D). We also analyzed the association of PSMA (*FOLH1)* expression with patient survival using the TCGA GBM dataset (n = 567 mRNA microarray samples). Consistent with our findings, no association with patient survival was observed when patients were separated based on the PSMA (*FOLH1)* expression as the highest 25% compared to the lowest 25% (*P* **=** 0.544, log-rank test) (Figure 2E) indicating that PSMA is not an independent prognostic factor in gliomas.

### 3.3 PSMA expression in the secondary brain tumors

Extending our study, we analyzed PSMA expression in the brain metastases (BMs) of breast and lung carcinomas and melanomas. Of the whole tissue sections analyzed, all 18/18 breast cancer BM samples (breast adenocarcinoma, n = 1, infiltrative duct carcinomas, n = 17) (Figure 3A,C) and 18/19 melanoma BMs (Figure 3B,D) expressed PSMA in their vasculature. Next, we compared PSMA expression in the primary lung tumors and their associated brain metastases (BM) and scored separately the vascular and tumor cell expression. The primary lung carcinoma samples in our tissue array were of various histological types containing i) non-small cell lung cancers such as adenocarcinomas and squamous cell carcinomas (SCC) and ii) small cell lung carcinomas (SCLC). PSMA was expressed in the vasculature of all types of lung carcinomas (Figure 3E,G,I) and their metastases (Figure 3F,H,J and Table 3). Overall the vascular expression of PSMA was highest in the lung cancer BMs (Figure 3F,H,J) and lowest in the melanoma BMs (Figure 3B, D) while the breast cancer BMs expressed moderate levels of PSMA (Figure. 3A,C) as judged by visual evaluation. We then compared PSMA expression in the primary lung carcinomas and their brain metastases from the same patients (n = 52). In addition to the vascular expression of PSMA, the primary lung carcinomas (Figure 4A,C) and their corresponding metastatic lesions (Figure 4B,D) also expressed PSMA in the tumor cells. PSMA expression in the vasculature of the primary lung carcinomas and their metastatic lesions had no significant association with various clinical factors (histopathological staging, age, or gender) (Table 4). Since smoking is one of the major causative factors in lung tumors, patients were divided into three categories based on their smoking history: i) no prior smoking, ii) stopped smoking, and iii) still a smoker. No association between smoking and either the vascular or the cellular expression of PSMA was detected in primary lung carcinomas or their BMs (Table 4).

**Table 3.** Histological classification of PSMA-positive tumors.

| ***Tumor*** |  | ***Tumor histology*** |  |  |  |  |
| --- | --- | --- | --- | --- | --- | --- |
|  |  | ***Adenocarcinoma*** | ***Neuroendocrine*** | ***SCC*** | ***SCLC*** | ***Others*** |
| Lung<br>carcinomas | Vascular | 47.5% | 50% | 57.1% | 50% | 50% |
|  |  | (19/40) | (1/2) | (4/7) | (2/4) | (1/2) |
|  | Tumor cells | 62.5% | 50% | 85.7% | 50% | 100% |
|  |  | (25/40) | (1/2) | (6/7) | (2/4) | (2/2) |
| Lung BMs | Vascular | 54.8% | 100% | 83.3% | 100% | 100% |
|  |  | (23/42) | (2/2) | (5/6) | (4/4) | (2/2) |
|  | Tumor cells | 71.4% | 100% | 100% | 100% | 100% |
|  |  | (30/42) | (2/2) | (6/6) | (4/4) | (2/2) |

**Table 4.** Clinicopathologic features of the primary lung carcinomas and their brain metastases.

| *Pathological parameter* |  | *Features* | *Primary lung carcinomas* |  |  |  | *Lung brain metastasis* |  |  |  |
| --- | --- | --- | --- | --- | --- | --- | --- | --- | --- | --- |
|  |  |  | *PSMA* |  | *Total* | *P-Value*<br>*Chi-square* | *PSMA* |  | *Total* | *P - Value*<br>*Chi-square* |
|  |  |  | *+* | *-* |  |  | *+* | *-* |  |  |
| **Pathological staging** | I-II | Vascular | 15 | 18 | 33 | 0.50 | 20 | 13 | 33 | 0.57 |
|  | III-IV |  | 11 | 9 | 20 |  | 13 | 6 | 19 |  |
|  | I-II | Tumor | 22 | 11 | 33 | 0.90 | 24 | 9 | 33 | 0.29 |
|  | III-IV | cells | 13 | 7 | 20 |  | 17 | 2 | 19 |  |
| **Gender** | Male | Vascular | 14 | 15 | 29 | 0.90 | 18 | 11 | 29 | 0.72 |
|  | Female |  | 13 | 13 | 26 |  | 18 | 9 | 27 |  |
|  | Male | Tumor | 20 | 9 | 29 | 0.56 | 22 | 7 | 29 | 0.61 |
|  | Female | cells | 16 | 10 | 26 |  | 22 | 5 | 27 |  |
| **Age** | Min | Vascular | 38 | 43 | Median | 0.57 | 38 | 43 | Median | 0.19 |
|  | Max |  | 77 | 71 | 60<sup>(†)</sup>/57<sup>(‡)</sup> |  | 73 | 71 | 58<sup>(†)</sup>/62<sup>(‡)</sup> |  |
|  | Min | Tumor | 38 | 43 | Median | 0.93 | 38 | 43 | Median | 0.13 |
|  | Max | cells | 77 | 71 | 59<sup>(†)</sup>/56<sup>(‡)</sup> |  | 73 | 71 | 58<sup>(†)</sup>/63<sup>(‡)</sup> |  |
| **Smoking** | 0-1<sup>(§)</sup> | Vascular | 10 | 10 | 20 |  | 11 | 5 | 16 | 0.83 |
|  | 2<sup>(¶)</sup> |  | 16 | 17 | 33 | 0.92 | 25 | 13 | 38 |  |
|  | 0-1<sup>(§)</sup> | Tumor | 12 | 8 | 20 |  | 12 | 4 | 16 | 0.71 |
|  | 2<sup>(¶)</sup> | cells | 22 | 11 | 33 | 0.62 | 31 | 7 | 38 |  |

(†)PSMA-positive
(‡) PSMA-negative
(§) Never smoked
(0) Quit smoking
(1) (¶) Still smoking (2)

**Figure 3.**
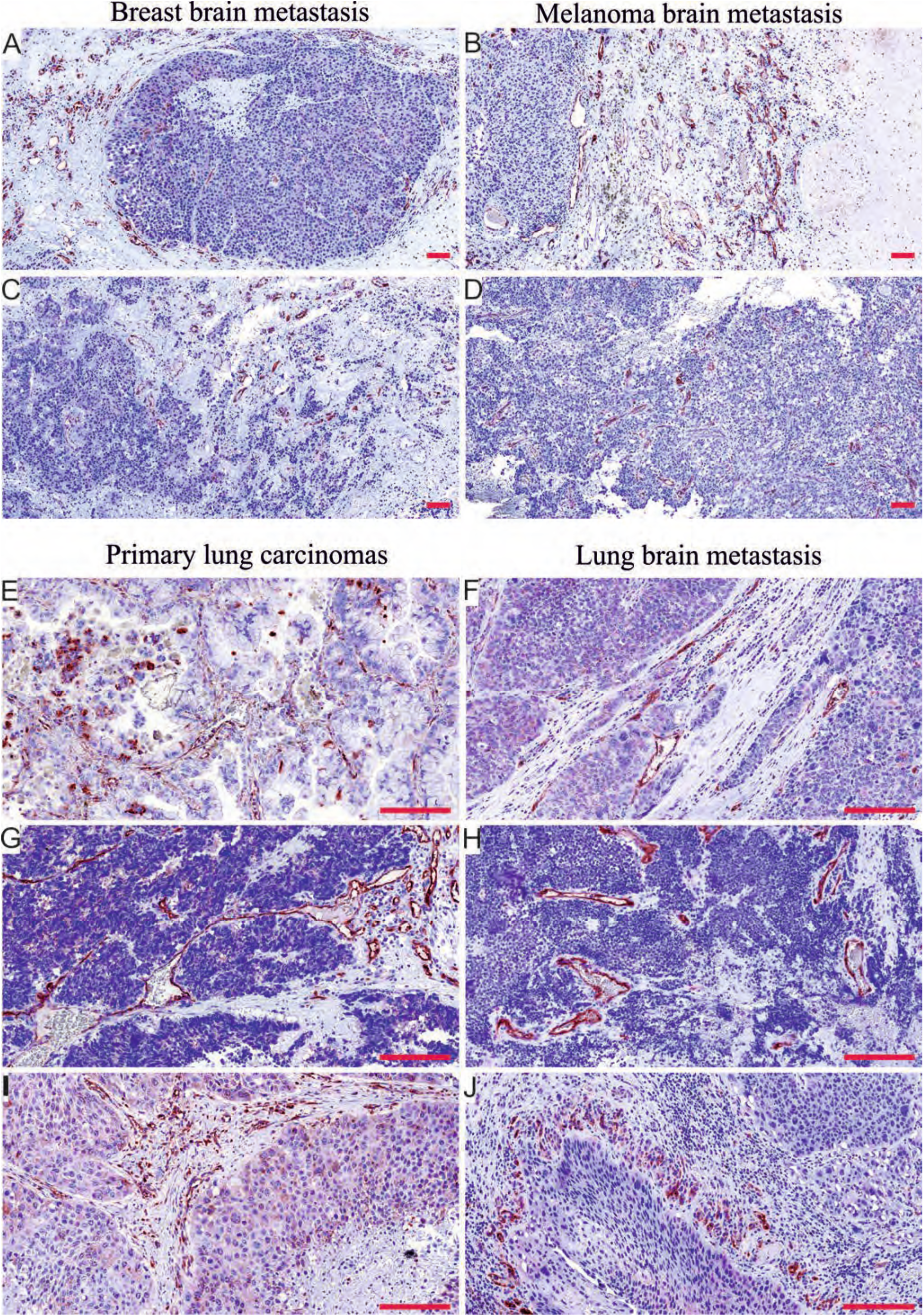
PSMA expression in the brain metastases of breast carcinomas and melanomas as well as in primary lung carcinomas and their associated metastases. A. PSMA is highly expressed in the vasculature of the brain metastases of breast adenocarcinoma and C. invasive ductal carcinoma as well as in B,D melanoma brain metastases. E,G,I. Vasculature of the primary lung carcinomas of different histological subtype such as adenocarcinoma (E), small cell lung carcinoma (G), and squamous cell carcinoma (I) were highly positivity for PSMA. F,H,J. The brain metastases of the lung adenocarcinoma (F), small cell lung carcinoma (H), and squamous cell carcinoma (J) also expressed PSMA in the vasculature. There was a pattern of cellular pseudopalisades (J) closely surrounded by the PSMA-positive vasculature similar to the one seen in glioblastomas. Scale bar = 100 μm.

**Figure 4.**
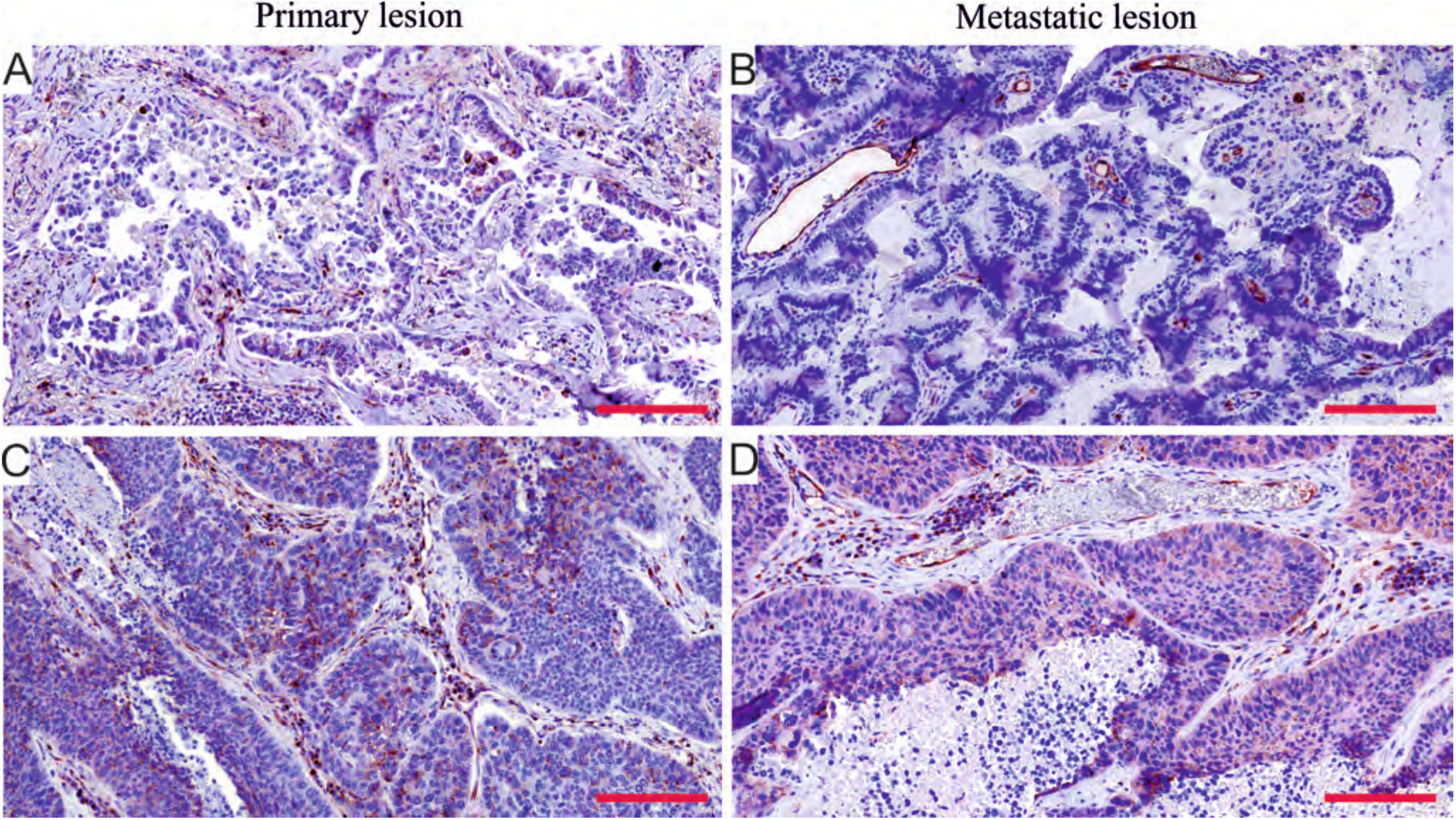
PSMA expression in the paired primary lung cancers and brain metastases from the same patient. A,B. PSMA expression in matched lung adenocarcinoma and brain metastasis. C,D. PSMA expression in matched lung squamous cell carcinoma and brain metastasis. Both the primary and metastatic tumor vasculature showed positivity for PSMA. Scale bar = 100 μm.

### 3.4 Tumor cell expression of PSMA is higher in secondary than in primary brain tumors

The tumor cell expression of PSMA varied substantially amongst the tumor types. When we compared PSMA expression between brain metastases of different origin (Figure 5 A-C) and gliomas (Figure 5D), PSMA expression in tumor cells was higher in the BMs of lung carcinoma than in the other BMs or in gliomas. The cellular expression of PSMA was very low in gliomas, which is consistent with the IvyGAP dataset results (Figure 2B). In addition, tumor cell expression of PSMA in the lung carcinomas and their metastatic lesions did not associate with any clinical or molecular pathological parameters (Table 4).

**Figure 5.**
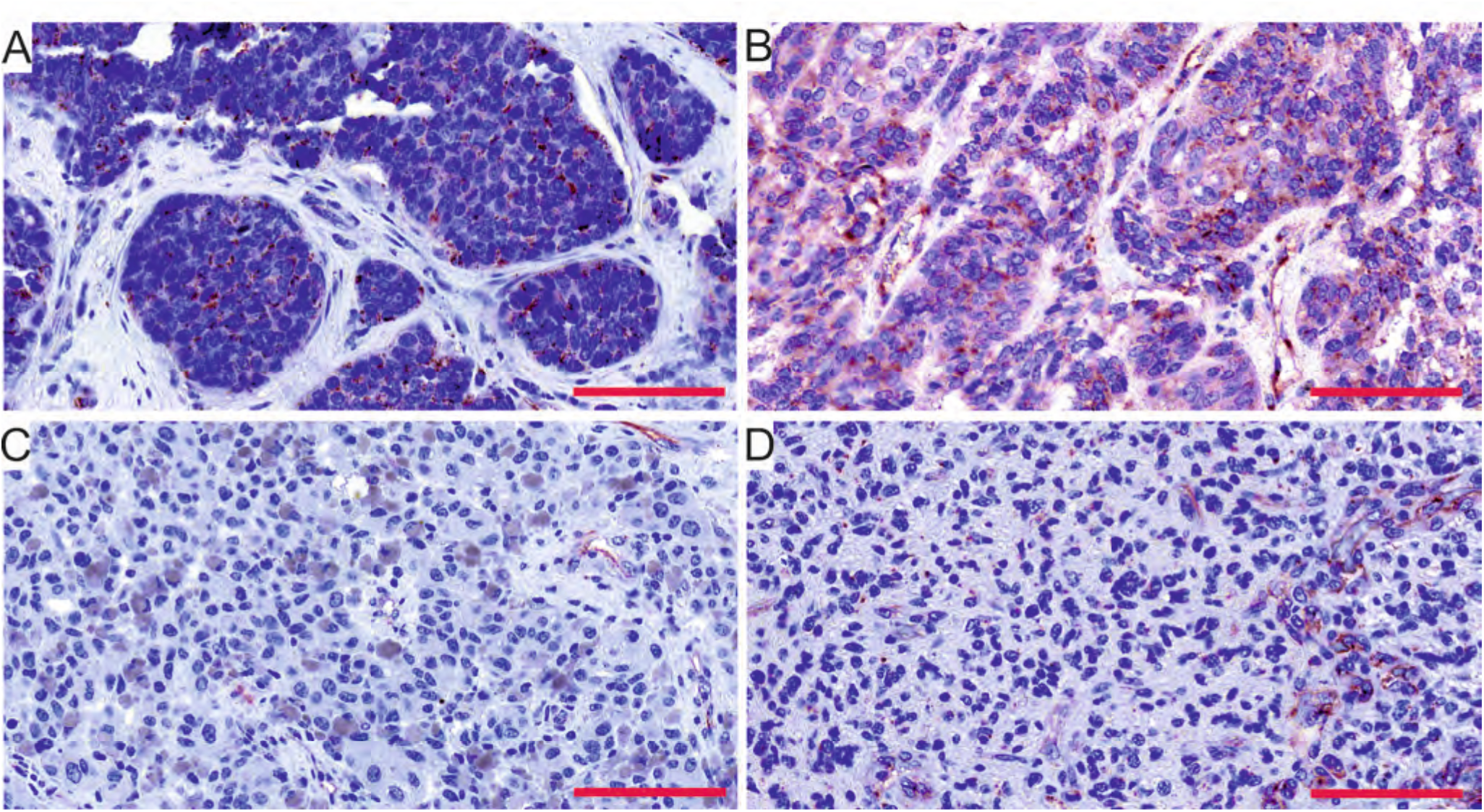
PSMA expression in the tumor cells varied between the tumor types. Amongst the primary and secondary brain tumors, the brain metastases of breast (A) and the lung (B) carcinomas showed the highest expression whereas the melanoma BMs (C) and glioblastomas (D) showed the lowest expression of PSMA in the tumor cells. Scale bar = 100 μm.

### 3.5 Vascular PSMA expression in primary lung carcinomas associated with significantly accelerated metastatic dissemination into brain

The overall survival data was known for 55 lung carcinoma patients. The vascular PSMA expression in the primary lung carcinomas associated with a tendency towards decreased overall survival (Mantel-cox *P* = 0.092 / Breslow *P* = 0.051) (Table 7, Figure 6A), whereas no association of PSMA expression in the tumor cells with the overall survival was observed (Mantel-cox *P* = 0.583/ Breslow *P* = 0.33) (Figure 6B). In addition, PSMA expression either in the tumor vasculature or in tumor cells did not associate with the progression-free survival (Table 5). Our patient cohort consisted of 52 matched primary lung carcinomas and their corresponding brain metastases. Primary tumors with PSMA positive vasculature associated with significantly faster metastatic dissemination to the brain compared to tumors with PSMA-negative vasculature (Mantel-cox *P* = 0.039 / Breslow *P* = 0.005) (Table 6, Figure 6C). No association between PSMA expression in the tumor cells and accelerated metastatic dissemination was observed (Mantel-cox *P* = 0.292 / Breslow *P* = 0.074) (Table 6, Figure S2A). The patients excluded from the analysis (n = 3) based on the diagnosis of the metastasis prior the primary tumor detection, still showed accelerated metastatic dissemination to the brain only if the vasculature was PSMA-positive (Mantel-cox *P* = 0.012 / Breslow *P* = 0.009) (Table S1, Figure S2B) but not the tumor cells (Mantel-cox *P* =0.135 / Breslow *P* = 0.046) (Table S1, Figure S2C).

**Table 5.**
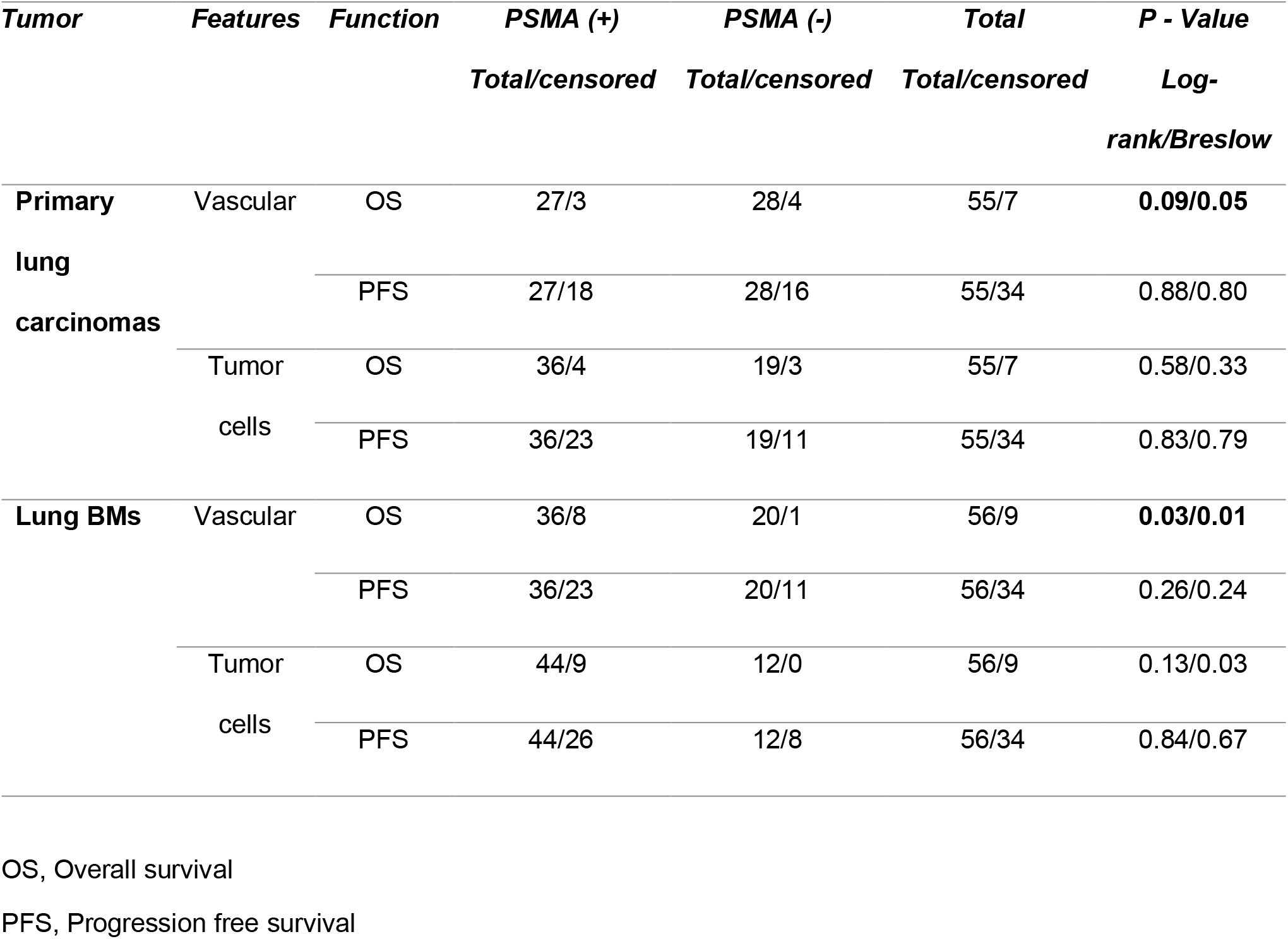
Overall survival and progression free survival.

**Table 6.**
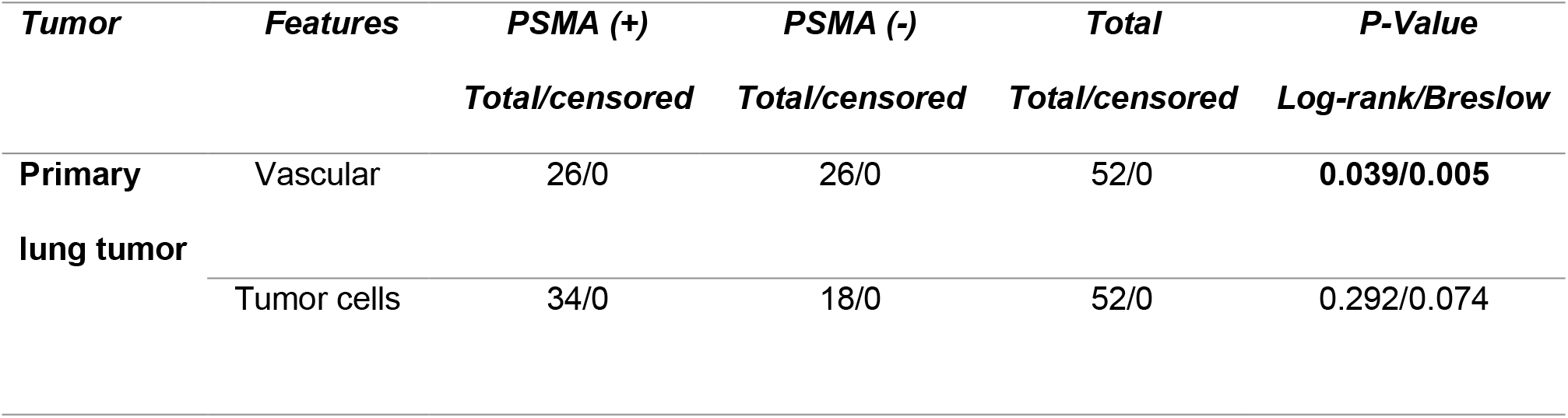
Time to metastatic dissemination from the primary tumor to the brain (n = 52).

**Figure 6.**
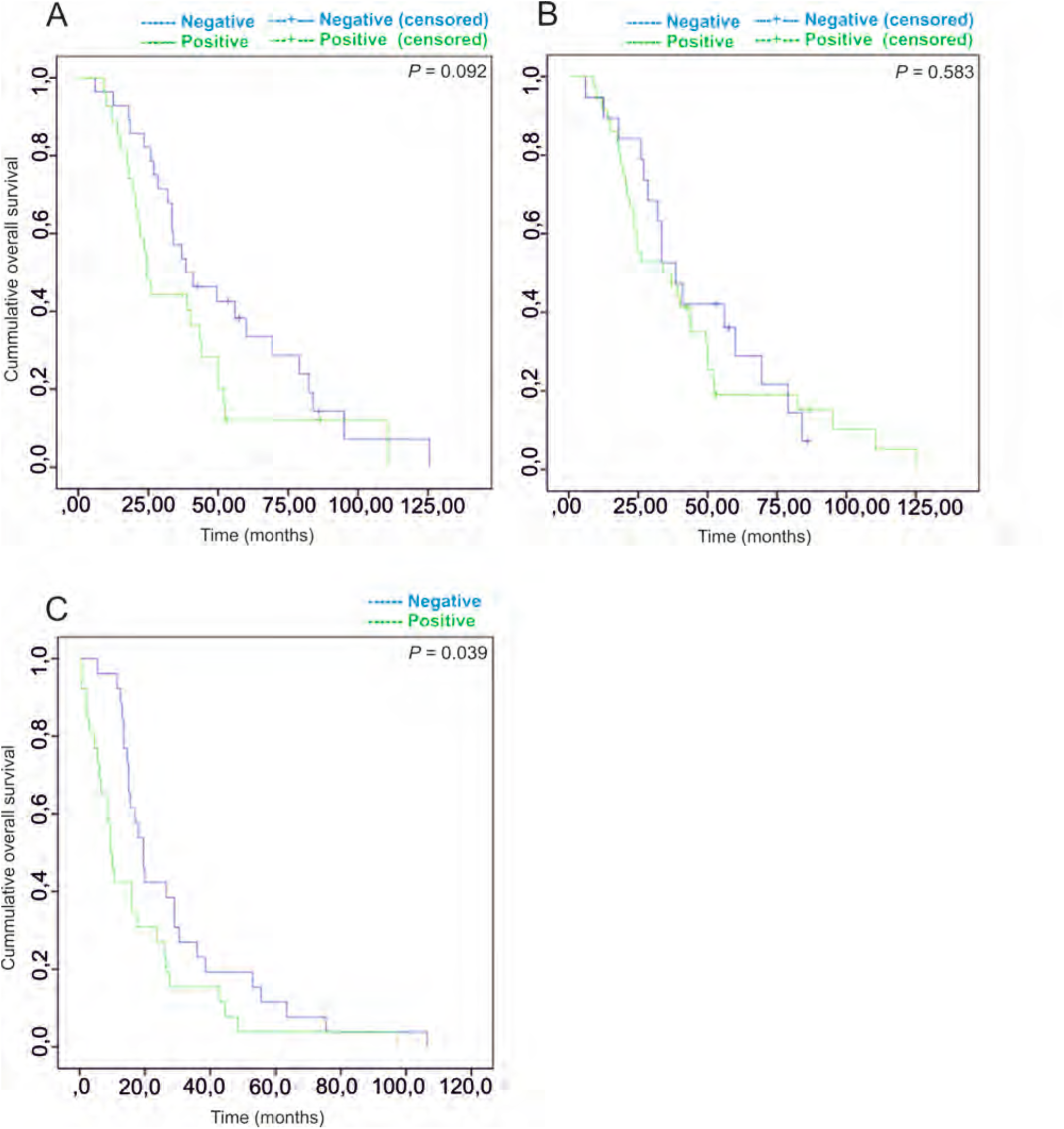
Association of PSMA expression with patient survival. A. The overall survival of the patients (n = 55) whose tumors expressed PSMA in the vasculature (Mantel-cox *P* = 0.092 / Breslow *P* = 0.051. B. The overall survival of the patients (n = 55) with PSMA expression in tumor cells (Mantel-cox *P* = 0.583 / Breslow *P* = 0.051). C. Primary lung carcinomas that showed vascular PSMA expression display accelerated rate of metastatic dissemination to brain (n= 52, Mantel-cox *P* = 0.039 / Breslow *P* = 0.005.

## 4. Discussion

In this report, we studied the expression of PSMA in large cohorts of primary and secondary brain tumors, using both traditional clinical and molecular pathological methods. PSMA expression has been reported previously in gliomas (15), breast BMs (20, 21) and in primary lung carcinomas (22) but with a very limited number of patients. Our study spans across diverse tumor types with a large set of patient samples including gliomas (n = 371) and brain metastases of tumors that commonly metastasize to the brain such as breast BMs (n = 18), melanoma BMs (n = 19) and lung BMs (n = 52, with matched primary lung carcinomas). We assessed PSMA expression at the protein level using tumor tissue microarray and associated it with various clinical and molecular pathological parameters following the WHO 2016 guidelines for the classification of Tumors of the Central Nervous System. We also took advantage of the cancer genome databases such as the TCGA (n = 567) and Ivy-GAP (n = 10) to assess PSMA expression at the RNA level.

Our results show that PSMA expression was highest in the vasculature of glioblastomas especially in the proliferating hyperplastic microvasculature, one of the key angiogenic and histopathological traits of glioblastomas. This result is in agreement with an earlier study that consisted of 14 glioma and 5 glioblastoma specimens (15). The study by Nomura *et al.* (15) showed moderate PSMA expression in grade I pilocytic astrocytomas (5 patients) and high expression in grade IV glioblastomas (GBM – 5 patients) while grade II diffuse (4 patients) and grade III anaplastic (5 patients) astrocytomas were negative. The discrepancy between our study and Nomura *et al.* (15) may be explained by the relatively small number of patients in each glioma grade in the Nomura study. They also pointed out that the variability of PSMA staining (both vascular and tumor cells) within gliomas needs to be explored in larger scale studies, which we have now addressed here.

Microvascular hyperplasia is characterized by tufted micro-aggregates, neovascular bodies with abundant and swiftly dividing endothelial cells, and inadequate pericyte/smooth muscle cell coverage. Microvascular hyperplasia, an essential feature associated with the tumor aggressiveness, is a potent predictor of poor prognosis (23). Our TMA-results corroborated with the results we obtained using the Ivy-GAP dataset. The significance of the Ivy-GAP project is the use of laser microdissection to sample individual anatomic structures of the tumor followed by RNA sequencing. This results in structure-specific gene expression signatures, unlike the conventional tumor RNA sequencing in which all different anatomical structures and cell types are pooled together. We also show that integrin β1 (*ITGB1*) is upregulated in the proliferating microvasculature. This is in concordance with the previous experimental report that showed PSMA-dependent increase in integrin β1-mediated signal transduction in a laminin-specific manner to regulate angiogenic endothelial adhesion and invasion (16).

There are ample studies utilizing PSMA-based imaging agents for tumor detection in prostate cancer (24-31), high-grade gliomas (32-36) and lung BMs (33), as well as in follicular thyroid adenoma (37), metastatic renal cell carcinoma (38) and in melanoma and small cell lung cancer (SCLC) xenografts *in vivo* (39). However, recent reports showed that the cerebral radionecrotic uptake (40) or stroke (41) resulted in false positive diagnoses of cerebral metastases based on PSMA/CT uptake as limitation of the PSMA-based diagnostic glioma imaging. A single arm phase II study where PSMA-recognizing ADC was used in progressive GBM patients (n = 6) following prior treatment with radiation, temozolomide, and bevacizumab showed no demonstrable activity in these patients (NCT01856933). This was likely due to the minimal expression of the PSMA target and associated with dose-limiting toxicity (42). In our study, the vascular expression of PSMA was observed in about 1/3 of glioblastomas. It thus, raises a question whether PSMA-based targeting agents could be used in these patients after patient stratification.

In addition to the glioma samples, we screened PSMA expression in the brain metastases of lung and breast cancers and melanomas. Even though PSMA expression has been reported earlier in breast cancer brain metastases (15, 33), our study shows the association of the vascular PSMA expression in primary lung tumors with the faster dissemination to the brain. A future study focusing on the mechanisms of how PSMA in the tumor endothelial cells accelerates metastatic dissemination to brain would provide important novel information. The tendency towards decreased overall survival of patients expressing PSMA in the vasculature of their lung tumors may be used as a prognostic factor but needs to be further validated in a larger data set of patient samples. In accordance with this, it has been reported that the vascular expression of PSMA in squamous cell carcinoma of the oral cavity is associated with poor prognosis (43).

PSMA expression in the proliferating microvasculature of glioblastoma near the cellular pseudopalisades is most likely a result of the hypoxia-induced angiogenesis. However, the vascular expression in primary lung neoplasm leading to accelerated metastatic dissemination has not been described earlier. Interestingly, a very recent preclinical study of a similar protein to PSMA, the prostate specific antigen (PSA), could provide some clues on the molecular mechanism responsible for the accelerated metastasis (44). In this study, the authors show how PSA activates the vascular endothelial growth factor C (via a proteolytic cleavage) which in turn promotes a pro-metastatic lymph/angiogenic niche.

In conclusion, we have addressed the under-explored PSMA/*FOLH1* expression in the primary and secondary brain tumors using large patient datasets as well as the genomic datasets. In the future, it will be interesting to see whether PSMA could be used as a diagnostic biomarker to detect and/or anticipate metastatic dissemination to the brain.

## Acknowledgments

This project received funding from the European Union’s Horizon 2020 research and innovation program under the Marie Skłodowska Curie grant agreement No-6420004, Jane and Aatos Erkko Foundation and Finnish Cancer Organizations.

The images were generated using the 3DHISTECH Panoramic 250 FLASH II digital slide scanner at Genome Biology Unit supported by the HiLIFE and the Faculty of Medicine, University of Helsinki, and Biocenter Finland. We thank CSC—IT Center for Science, Ltd. (https://www.csc.fi/csc) for providing the computational resources. The results published here are in part based upon data generated by The Cancer Genome Atlas project (dbGaP Study Accession: phs000178.v9.p8) established by the NCI and NHGRI. Information about TCGA and the investigators and institutions who constitute the TCGA research network can be found at http://cancergenome.nih.gov.

## Authors’ contributions

Study conceived, designed and manuscript written by: JTR, VLJ and PL

Experiments performed /computational study conceived by: JTR

Computational study conceived and performed by: SL

Statistical analysis performed by: HH, HS

Sample collection and pathological analysis: JT, HH, LR, VT, JM

Study supervised by: JT, MN, VLJ and PL

## Informed consent

Patients included in the study have provided the informed consent.

